# Secondary contact and local adaptation contribute to genome-wide patterns of clinal variation in *Drosophila melanogaster*

**DOI:** 10.1101/009084

**Authors:** Alan O. Bergland, Ray Tobler, Josefa González, Paul Schmidt, Dmitri Petrov

## Abstract

Populations arrayed along broad latitudinal gradients often show patterns of clinal variation in phenotype and genotype. Such population differentiation can be generated and maintained by historical demographic events and local adaptation. These evolutionary forces are not mutually exclusive and, moreover, can in some cases produce nearly identical patterns of genetic differentiation among populations. Here, we investigate the evolutionary forces that generated and maintain clinal variation genome-wide among populations of *Drosophila melanogaster* sampled in North America and Australia. We contrast patterns of clinal variation in these continents with patterns of differentiation among ancestral European and African populations. Using established and novel methods we derive here, we show that recently derived North America and Australia populations were likely founded by both European and African lineages and that this admixture event contributed to genome-wide patterns of parallel clinal variation. The pervasive effects of admixture meant that only a handful of loci could be attributed to the operation of spatially varying selection using an *F*_*ST*_ outlier approach. Our results provide novel insight into the well-studied system of clinal differentiation in *D. melanogaster* and provide a context for future studies seeking to identify loci contributing to local adaptation in a wide variety of organisms, including other invasive species as well as some temperate endemics.

## Introduction

All species live in environments that vary through time and space. In many circumstances, such environmental heterogeneity can act as a strong selective force driving adaptive differentiation among populations. Thus, a major goal of evolutionary and ecological genetics has been to quantify the magnitude of adaptive differentiation among populations and to identify loci underlying adaptive differentiation in response to ecologically relevant environmental variation.

Phenotypic and genetic differentiation between populations has been examined in a variety of species. In some cases, patterns of differentiation are directly interpretable in the context of circumscribed environmental differences that occur over short spatial scales (Richardson et al. 2014) For instance, differences in salinity experienced by freshwater and marine populations of sticklebacks has led to the identification of key morphological, physiological, and genetic differences between replicate pairs of populations (Kitano et al. 2010; Jones et al. 2012). Similarly, pigmentation morph frequency closely tracks variation in substrate color for a variety of species (reviewed in Gray & McKinnon 2007) thereby providing an excellent opportunity to directly relate environmental variation to phenotypic and genetic differentiation.

Patterns of genetic and phenotypic variation have also been examined in species arrayed along broad geographical transects such as latitudinal clines (Endler 1993). In this paradigm, the goal has often been to identify the phenotypic and genetic basis for adaptation to temperate environments. In certain cases it has been possible to directly relate latitudinal variation in specific environmental variables to aspects of phenotypic and genetic differentiation (e.g., photoperiod and critical photoperiod or flowering time; Bradshaw & Lounibos 1977; Stinchcombe *et al.* 2004). In general, the collinearity of multiple ecological and environmental variables along latitudinal clines often complicates the direct relation of environmental variation to specific phenotypic and genetic differences. Nonetheless, because many genetically based phenotypic clines within species often mirror deeper phylogenetic differentiation between endemic temperate and tropical species, it is clear that populations distributed along latitudinal clines have adapted to aspects of temperate environments (Gibert et al. 2001).

Latitudinal clines have been extensively studied in various drosophilid species, most notably *Drosophila melanogaster.* Parallel clines in morphological (Rajpurohit et al. 2008; Telonis Scott et al. 2011), stress tolerance (Hoffmann et al. 2002; Schmidt et al. 2005), and life-history traits (Schmidt et al. 2005; Lee et al. 2011) have been identified in *D. melanogaster* populations distributed along multiple continents. These phenotypic clines demonstrate that flies from poleward locales are generally more hardy albeit less fecund, reflecting a classic trade-off between somatic maintenance and reproductive output (Schmidt et al. 2005) that would be alternately favored between populations exposed to harsh winters versus more benign tropical environments. Extensive clinal variation in various genetic markers has also been identified (Sezgin *et al.* 2004; Hoffmann & Weeks 2007). In some cases clinal genetic variants have been directly linked to clinally varying phenotypes (Schmidt *et al.* 2008; Paaby *et al.* 2010; Lee *et al.* 2013; Paaby *et al.* 2014a), whereas in other cases parallel clinal variation at genetic markers has been documented across multiple continents (Turner et al. 2008; González et al. 2010; Reinhardt et al. 2014). Taken as a whole, there is abundant evidence that local adaptation to spatially varying selection pressures associated with temperate environments has shaped clinal patterns of phenotypic and genetic variation in *D. melanogaster*.

Demographic forces can also heavily shape patterns of clinal variation (Endler 1993) and the recent demographic history of *D. melanogaster* may be particularly relevant to our understanding of clinal patters of genetic variation in this species. *D. melanogaster* is an Afro-tropical species (David & Capy 1988) that has colonized the world in the wake of human migration. Population genetic inference suggests that *D. melanogaster* first migrated out of Africa to Eurasia approximately 15,000 years ago (Li & Stephan 2006) and eventually migrated eastward across Asia, arriving to South East Asia approximately 2.5Kya (Laurent et al. 2011). *D. melanogaster* invaded the Americas and Australia within the last several hundred years and likely colonized these continents in their entirety quickly (Bock & Parsons 1981; Keller 2007). Historical records suggest that *D. melanogaster* colonized North America and Australia each in a single event (Bock & Parsons 1981; Keller 2007).

In contrast, however, population genetic (Caracristi & Schlötterer 2003; Duchen *et al.* 2013; Kao *et al.* 2014) and morphological evidence (Capy et al. 1986; Ferveur et al. 1996; Coyne et al. 1999; Takahashi et al. 2001; Rouault et al. 2004) suggest that, for the Americas at least, there were multiple colonization events with some migrants coming from Africa and some from Europe. While there is less evidence that Australia experienced multiple waves of colonization by *D. melanogaster* such a scenario is plausible given the high rates of human migration and inter-continental travel during the 19^th^ century. If North America and Australia experienced multiple waves of immigration from highly differentiated ancestral populations (Pool et al. 2012), many genetic variants would appear clinal, even in the absence of spatially varying selection pressures (Endler 1993; Caracristi & Schlötterer 2003).

Investigating whether North America and Australia represent secondary contact zones is, therefore, crucial for our understanding of the extent of spatially varying selection operating on this species. We note that the models of dual colonization and adaptive differentiation as evolutionary forces that generate and maintain clinal variation in North America and Australia are not mutually exclusive. Notably, one plausible model is that dual colonization of these continents generated patterns of clinal variation and spatially varying selection has subsequently slowed the rate of genetic homogenization among populations. Accordingly, we sought to investigate whether genome-wide patterns of clinal genetic variation in North America and Australia show signals of dual colonization and local adaptation.

We find that both North American and Australian populations show several genomic signatures consistent with secondary contact and suggest that this demographic process is likely to have generated patterns of clinal variation at a large fraction of the genome in both continents. Despite this genome-wide signal of recent admixture, we find evidence that spatially varying selection has shaped patterns of allele frequencies at some loci along latitudinal clines using an *F*_*ST*_ outlier approach. Some of these *F*_*ST*_ outliers are in or near genes previously identified to underlie life-history and stress tolerance traits. However, in other cases, previously identified functional polymorphisms were not identified as *F*_*ST*_ outliers demonstrating the inherent challenge of identifying the genetic targets of spatially varying selection. We discuss these findings in relation to the well-documented evidence of spatially varying selection acting on this species as well as the interpretation of patterns of genomic variation along broad latitudinal clines in general.

## Materials and Methods

### Genome-wide allele frequency estimates

We utilized novel and publically available genome-wide estimates of allele frequencies of *D. melanogaster* populations sampled world-wide (Figure 1, Table S1). Allele frequency estimates of six North American populations are described in Bergland (2014). Allele frequency estimates of three European populations are described in Bastide et al. (2013) and Tobler et al. (2014). Allele frequency estimates from 20 African populations with full genome sequence that are described in Pool et al. (2012). Allele frequency estimates of two Australian populations are described in Kolaczkowski et al. (2011). Allele frequency estimates from an additional two Australian populations are reported here for the first time. Allele frequency estimates from these additional Australian populations were made by pooling ten individuals from each of 22 isofemale lines originating from Innisfail (17°S) or Yering Station (37°S), Australia (isofemale lines kindly provided by A. Hoffmann). Sequencing library preparation and mapping followed methods outlined in Bergland et al. (2014). Because Australian data were low coverage (~10X per sample, on average), we combined the two northern populations and two southern populations into two new, synthetic populations which we refer to as ‘tropical’ and ‘temperate,’ respectively. In several analyses, *D. simulans* was used as an outgroup. In these analyses, we used genome-wide allele frequency estimates from a world-wide collection *D. simulans* (Begun et al. 2007), using genome-wide *lift-over* files reported in Bergland et al. (2014).

**Figure 1.**
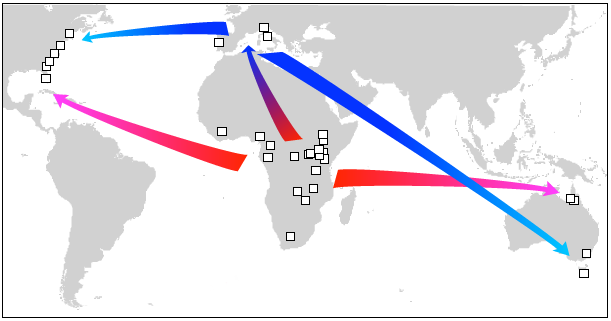
Map of collection locales (squares) and proposed colonization routes of *D. melanogaster* (arrows).

We performed SNP quality filtering similar to the methods presented in Bergland et al. (2014). Briefly, we excluded SNPs within 5bp of polymorphic indels, SNPs within repetitive regions, SNPs with average minor allele frequency less than 15% in both North America and Australia, SNPs with low (<5) or excessively high read depth (>2 times median read depth) and SNPs not present in the Drosophila Genetic Reference Panel (Mackay et al. 2012). African samples were not quality filtered for read depth because allele frequency estimates from these samples were derived from sequenced haplotypes and not pooled samples. Regions of inferred admixture (Pool et al. 2012) in African samples (i.e., introgression of European haplotypes back to African populations) were removed from analysis.

### Estimation of the population tree

We calculated Nei’s genetic distance (Nei 1972) between each pair of populations as,

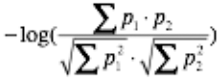

where *p_1_* and *p_2_* are allele frequency estimates in population 1 and 2, and generated a population tree using the neighbor-joining algorithm implemented in the R (R Core Team 2014) package *ape* (Paradis et al. 2004). To generate bootstrap values for each node, we randomly sampled 10,000 SNPs 100 times. We generated estimates of the population tree using the whole genome and for each chromosome focusing on SNPs that occur within the large, cosmopolitan inversions (Corbett-Detig & Hartl 2012), outside these inversions, or both.

### Estimation of the proportion African and European ancestry

We used a simple linear regression method to estimate the proportion of African and European ancestry for each newly derived North American and Australian population. This method follows an approach outlined in Alkorta-Aranburu et al. (2012) where each newly derived population is modeled as a linear combination one African and one European population using an intercept-free regression model. Ancestry coefficients were calculated averaging over 100 bootstrap replicates of 5000 SNPs per derived population per pair of ancestral populations. We generated ancestry estimates using the whole genome and for each chromosome separately focusing on SNPs that occur within the large, cosmopolitan inversions, outside these inversions, or both.

To verify that our regression based method of ancestry proportion estimation is accurate and robust to various demographic scenarios, we simulated several demographic models using coalescent simulations as implemented in *ms* (Hudson 2002). To assess the accuracy of estimated ancestry proportion, we first simulated a three population model where with one ancestral population, one ancient population that diverged from the ancestral population 0.1*N*_*e*_ generations ago, and one derived population that results from admixture between the ancestral and ancient populations 0.001*N*_*e*_ generations ago. In these simulations, we varied the proportion of lineages in the derived population that originated from the ancient and ancestral populations. A graphical model of this demographic history is presented in Supplemental Figure 2 and *ms* code is available from DataDryad under accession doi: 10.5061/dryad.gg5nv.

**Figure 2.**
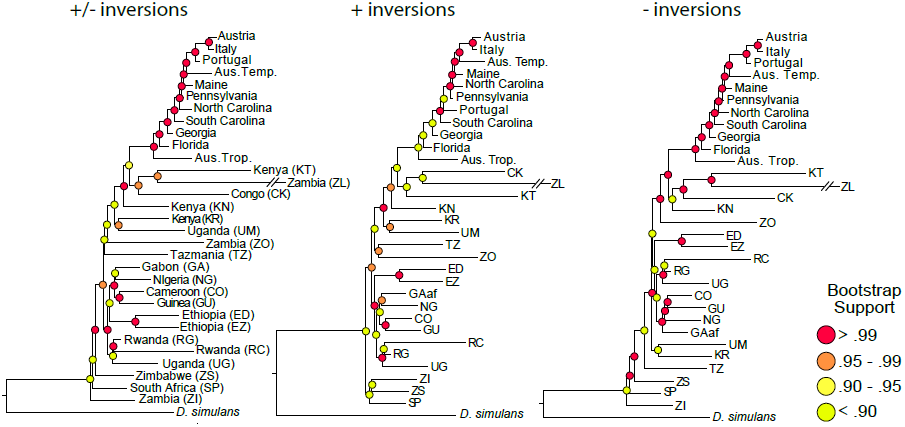
Estimated population tree of sampled locales. Loci were sampled across all chromosomes, focusing on SNPs residing within (+), outside (-) large, cosmopolitan inversions, or both(±). See Supplemental Figure 1 for population trees based on each chromosome.

Next, we sought to verify that single- and dual-colonization scenarios produce distinctive ancestry proportion clines. In these models, we simulated an ancient population that diverged from an ancestral population *0.1Ne* generations ago. In the single-colonization scenario a newly derived population was founded from the ancestral population and a series of four additional populations were derived from this initial newly derived population through a series of serial founder events. In the dual-colonization scenario, two newly derived populations were founded by either the ancestral or the ancient population. These newly derived populations each gave rise to another newly derived population though a serial founder event, and a fifth newly derived population resulted from the merger of these later populations. In both single- and dual-colonization models, migration was only allowed between neighboring newly derived populations. A graphical representations of these models can be found in Supplemental Figures 3 and *ms* code is available from DataDryad under accession doi:10.5061/dryad.gg5nv. In the single- and dual-colonization models, we varied the age of initial colonization of the newly derived populations and the extent of the population bottleneck during the initial and serial founder event(s).

**Figure 3.**
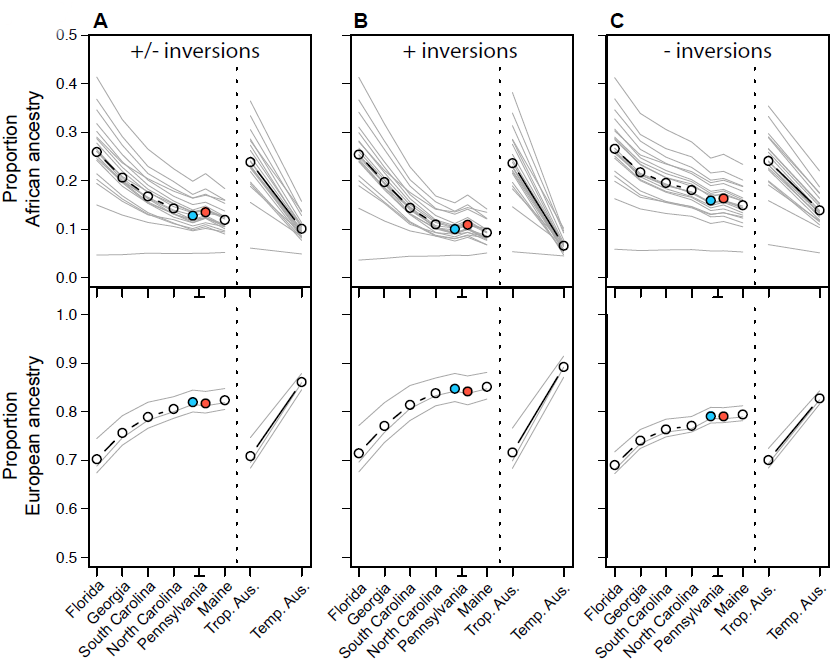
Proportion African and European ancestry in North American and Australian populations. Thin grey lines represent the ancestry estimate for any particular African or European population averaged over all European and African populations. Black line represents the average ancestry proportions from all African and European populations. The blue circle represents the spring Pennsylvania sample and the red circle represents the fall Pennsylvanian sample. See Supplemental Table X for ancestry estimates for each pairwise combination of African and European population, and for SNPs on each chromosome, and with locations inside and/or outside inversions.

### Formal tests of admixture

To further assess whether newly derived North America and Australian populations result from admixture of European and African lineages of flies, we performed formal tests of admixture using the *f*_3_ and *D* statistics (Reich et al. 2009; Patterson et al. 2012). Briefly, these statistics assess whether a proposed tree topology is consistent with the data.

The *f*_3_ statistic, denoted (C; A, B) assesses whether the data are consistent with the topology ((A, C), B) or ((B, C), A). A significantly negative *f_3_* statistic demonstrates that the data are consistent with both topologies, thus indicating that population C is derived from an admixture event from populations A and B or populations closely related to A and B. However, non-significant or positive *f_3_* statistic does not preclude the possibility of admixture (Patterson et al. 2012).

The *D* statistic, which is derived and conceptually similar to the ABBA-BABA statistic (Patterson et al. 2012), assesses whether the data are consistent with the proposed topology ((W, X), (Y, Z)). In general, a significantly positive *D* statistic indicates that population W results from admixture between population X and Y, or populations closely related to X and Y. A significantly negative *D* statistic can indicate that population X results from admixture between population W and Y, or populations closely related to W and Y. While, in general, the sign of significant *D* statistics can be difficult to interpret the most general interpretation is that the data do not conform to the proposed ((W, X), (Y, Z)) topology.

There are three possible *D* statistics any four populations: ((W, X), (Y, Z)), ((W, Y), (X, Z)), and ((W, Z), (X, Y)). In our analysis, we used *D. simulans* as an outgroup (population Z in this notation). Accordingly, the first two *D* statistics are most easily interpretable in our analysis. In this context, population W represents the newly derived North American or Australian population and population X and Y represent the putative source African and European populations, respectively. To simplify the interpretation of these *D* statistics, and to provide a conservative analysis of admixture, we report the *D* statistic corresponding to the minimum absolute *D* of ((W, X), (Y, Z)) and ((W, Y), (X, Z))

For each North American and Australian population, we calculated *f_3_* and *D* using each European population as one putative donor population and each African population as the other putative donor population. *f_3_* and *D* statistics were calculated using the *threepop* and *fourpop* programs included in *TreeMix* version 1.13 (Pickrell & Pritchard 2012) with 500 bootstrap replicates using a block size of 500 SNPs. Multiple testing correction was performed using the Bonferroni correction applied across all *p*-values from *f_3_* or *D* statistics.

### *F*_*ST*_ outlier identification

We used the *T*_*F-LK*_ statistic (Bonhomme et al. 2010) to identify *F*_*ST*_ outliers in North America and Australia. This statistic is related to classic Lewontin-Krakauer test for *F*_*ST*_ outliers, under the assumption that the distribution of *F*_*ST*_ is proportional to a *χ*^2^ distribution with degrees of freedom equal to one less the number of populations examined (Lewontin & Krakauer 1973). Various assumption underlying Lewontin-Krakauer test have been criticized (Robertson 1975) and the *T*_*F-LK*_ statistic attempts to correct for these by conditioning the distribution of *F*_*ST*_ values on the inferred underlying population tree. Under a variety of demographic scenarios (Bonhomme et al. 2010; Mita et al. 2013), including secondary contact (Lotterhos & Whitlock 2014), the *T*_*F-LK*_ statistic has generally been found to have low false positive-and high true positive-rates in *F*_*ST*_ outlier detection. Multiple testing correction for the *T*_*F-LK*_ statistic was performed using the false discovery rate methods implemented in the *q*-value package (Storey & Tibshirani 2003).

### Differentiation and rates of parallelism at various SNP classes

To assess rates of co-differentiation we calculated the odds that SNPs fell above one of three *F*_*ST*_ quantile thresholds (85, 90, 95%) in both North America and Australia. We compared this value to the odds of co-differentiation from 500 sets of randomly selected SNPs that were matched to the focal SNPs by recombination rate (Comeron et al. 2012), chromosome, inversion status (at the large, cosmopolitan inversions *In(2L)t, In(2R)NS, In(3L)Payne, In(3R)K, In(3R)Payne, In(3R)Mo, In(X)A,* and *In(X)Be)*, average read depth in North America and Australia, and heterozygosity in both continents. To control for the possible autocorrelation in signal along the chromosome, we divided the genome into non-overlapping 50Kb blocks and randomly sampled, with replacement, one SNP per block 500 times.

Next, we tested if SNPs at various annotation classes (e.g., short-introns, synonymous, non-synonymous, UTR; Figure 7) were more likely than expected by chance to be co-differentiated or show parallel changes in allele frequency between temperate and tropical locales in both North America and Australia conditional on them being co-differentiated. To assess rates of parallelism, we calculated the fraction of SNPs that were significantly co-differentiated and varied in a parallel fashion between North America and Australia for each SNP class and their matched, genomic controls, again controlling for the spatial distribution of SNPs along the chromosome. We report the difference in rates of parallelism. Standard deviations of the log_2_(odds-ratio) of co-differentiation and for differences in the rates of parallelism are calculated as in Bergland et al. (2014).

## Results

### Data

We examined genome-wide estimates of allele frequencies from ∼30 populations of *D. melanogaster* sampled throughout North America, Australia, Europe and Africa (Fig. 1). Our analyses largely focused on patterns of variation in North American and Australian populations and, consequently, we primarily focus on two sets of SNP markers. First, we utilized allele frequency estimates at ∼500,000 high quality SNPs that segregate at intermediate frequency (MAF > 15%) in North America. The second set was composed of ∼300,000 SNPs that segregate at intermediate frequency in Australia. For analyses that examine patterns of polymorphism in both North America and Australia, we examined SNPs that were at intermediate frequency in both continents, yielding a dataset of ∼190,000 SNPs. Because of the low sequencing coverage in the Australian populations, it is unclear if the reduced polymorphism in that continent reflects the demographic history of those populations or experimental artifact. Although our analysis primarily focused on patterns of polymorphism in North America and Australia, we also examined allele frequency estimates at both sets of polymorphic SNPs in populations sampled in Europe and Africa.

### Genomic signals of secondary contact

We performed a series of independent analyses to examine whether North America and Australia represent secondary contact zones of European and African populations of *D. melanogaster*. First, we constructed a neighbor-joining tree based on genome-wide allele frequency estimates from populations sampled world-wide using 100 sets of 10,000 randomly sampled SNPs. Neighbor joining trees were generated for SNPs residing within inversions, outside inversions, or both genome-wide (Figure 2) and for each chromosome separately (Supplemental Text 1). As expected, African populations exhibited the greatest diversity (Pool et al. 2012) and clustered at the base of the tree while European populations clustered at the tip. North American and Australian populations generally clustered between African and European populations (Figure 2), a pattern that supports the model (Kopelman et al. 2013) that both North American and Australian populations result from secondary contact of European and African ones.

In general, the topology of these population trees was independent of which SNPs were included in the analysis (i.e., compare Figure 2 and Supplemental Figure 1). One notable exception was the placement of the Portuguese population which falls within the North American samples when examining SNPs within inversions. In addition, the topology of the African populations varied depending on the set of SNPs under consideration. However the bootstrap support for the nodes among African populations are generally low, likely stemming from low coverage and imprecise estimates of allele frequencies in these populations. In chromosome specific analyses (Supplemental Text 1), nodes that do not agree with the genome-wide analysis generally had low boot-strap support.

Next, we calculated the proportion of African ancestry in North American and Australian populations by modeling these populations as a linear combination of African and European ancestry. Neutral coalescent simulations using *ms* found that this method accurately estimates ancestry proportions (Supplemental Figure 2). These simulations also revealed that a cline in ancestry proportion is unlikely under a single colonization scenario, even with extensive drift (Supplemental Figure 3), yet is stable and persistent under a dual-colonization scenario for long periods of time following initial colonization (Supplemental Figure 3).

In general, the proportion of African ancestry in North American and Australian populations is negatively correlated with latitude (Figure 3). Conversely, the proportion of European ancestry is positively correlated with latitude (Figure 3). One notable exception to these general ancestry patterns is a single population from Zambia (population ZI from Pool et al. 2012). This population is notable in that it has an exceptionally long branch in our neighbor joining analysis (Figure 2), likely indicating that it may be highly diverged from other African populations. In addition, the X-chromosome lacks a significant pattern of clinal variation in ancestry (Supplemental Figure 4), a pattern consistent with analyses presented in Kao et al. (2014).

Nonetheless, the general pattern of decreasing African ancestry with increasing latitude remains when using nearly any combination of African and European populations and for SNPs inside inversions, outside inversions, or both sampled genome-wide or on any particular autosome. Moreover, our estimate of ancestry proportion of the North Carolina population is in agreement with estimates made using other methods and data-sets (Duchen et al. 2013; Kao et al. 2014).

Finally, we calculated *f_3_* and *D* statistics (Reich et al. 2009; Patterson et al. 2012) – commonly referred to as formal tests of admixture – for each North American and Australian population using each sampled European and African population as a putative source population. We observe significantly negative *f_3_* statistics (Figure 4A) for each North American population when using the Italian or Austrian populations as European source populations and various African populations as the alternate source population (Supplemental Table 1). Significant admixture from Portuguese population into North America was only observed in southern populations using the f_3_ statistic. In general, *f_3_* statistics were not significantly negative for the Australian populations after correcting for multiple testing. Note, however, that temperate and tropical populations in Australia show evidence of admixture using the *f_3_* statistic before stringent Bonferroni multiple testing correction and remain marginally significant following multiple testing correction (Supplemental Table 1).

**Figure 4.**
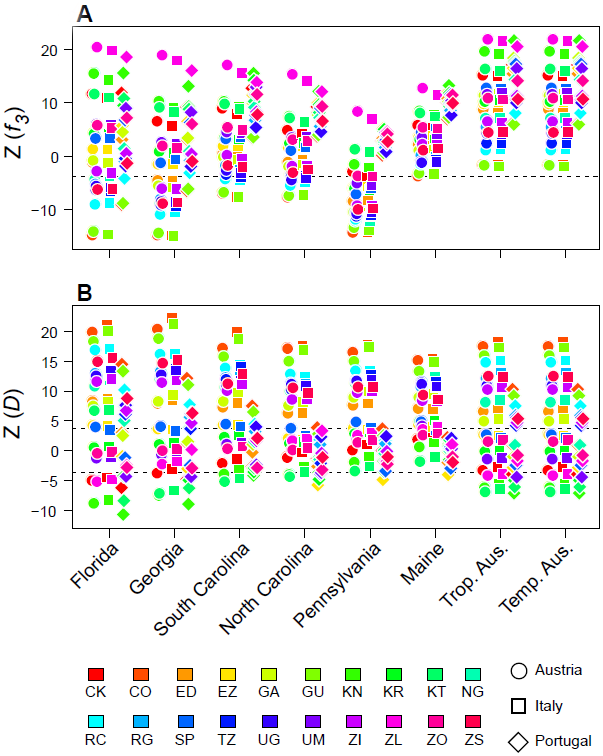
Observed standardized *Z* scored of (A) *f*_3_ and (B) *D* statistics for each North American and Australian population when considering every possible combination of African and European population and a putative donor population. The dashed line represents the Bonferroni significance threshold at alpha < 0.05. Values below the threshold in (A) are significantly different from 0. Values above or below the threshold in (B) are significantly different from 0.

We observe significantly positive *D* statistics for each newly derived North American and Australian population using various combinations of European and African populations as putative source populations (Figure 4B, Supplemental Table 1). Similar to the *f_3_* analysis, the Portuguese population shows little evidence of admixture into populations sampled in northern North America. It is worth noting that evidence of admixture using the *f_3_* and *D* statistics does not conclusively demonstrate that the sampled donor populations (i.e., the European and African populations) are the actual donor populations. Rather, evidence of admixture using these statistics implies that the sampled donor populations, or other unsampled yet closely related populations, are likely the donor populations.

Taken together, these results support the view that both North America and Australia represent secondary contact zones between European and African lineages of *D. melanogaster.* Our results confirm an earlier model (David & Capy 1988) and recent genomic evidence (Kao et al. 2014) that European *D. melanogaster* colonized high latitude locales in North American and Australia whereas African flies colonized low latitude locales in these regions. Genome-wide, low-latitude populations are more similar to African ones whereas high-latitude populations are more similar to European ones.

Under this dual-colonization scenario, we would expect that a large fraction of the genome varies clinally. Indeed, among North American populations of *D. melanogaster* approximately one third of all common SNPs, on the order of 10^5^, are clinal (following the analysis in Bergland et al. 2014). The vast extent of clinal variation in North America, then, is consistent with a dual colonization scenario which would generate patterns of clinal variation at a large fraction of the genome. However, these results do not preclude the existence of spatially varying selection that could also be acting among these populations which could explain patterns of differentiation reported for some loci (e.g., (Sezgin *et al.* 2004; Hoffmann & Weeks 2007; Paaby *et al.* 2010; 2014)) and could slow the rate of homogenization of allele frequencies at neutral polymorphisms throughout the genome among clinally distributed populations. We note that a similar analysis of the extent of clinality in Australia is not possible because we lack genome-wide allele frequency estimates from intermediate latitude populations in that continent.

### Genomic signals of parallel local adaptation along latitudinal gradients

Our previous analysis supports the model that the demographic history of *D. melanogaster* has contributed to genome-wide patterns of differentiation among temperate and tropical populations of *D. melanogaster* living in North America and Australia. Regardless of this putative demographic history, multiple lines of evidence suggest that populations of flies living along broad latitudinal gradients have adapted to local environmental conditions that may be associated with aspects of temperate environments (see *Discussion*). Accordingly, we performed several tests to assess whether there is a strong, observable genomic signal of local adaptation.

First, we sought to identify *F*_*ST*_ outliers using the *T*_*F-LK*_ method that attempts to identify SNPs subject to spatially varying selection while maintaining a high power and low false positive rate (Bonhomme et al. 2010; Mita et al. 2013; Lotterhos & Whitlock 2014). This method models the distribution of *F*_*ST*_ values after conditioning on the observed population tree among the sampled populations. We identified several hundred significantly differentiated SNPs in North America (Figure 5; Supplemental Table 2), some of which are in or near genes that have been previously implicated in adaptation to spatially varying selection pressures (e.g., *Abd-B;* Fabian et al. 2012) or likely affect life-history traits and correlates (e.g., *AlstR, TyR, DopR, sNPF*) via modulation of endocrine signaling (Bergland 2011). Intriguingly however, the amino acid polymorphism in *cpo* (3R: 13793588) previously implicated in clinal variation in diapause propensity (Schmidt et al. 2008) is significantly differentiated prior to, but not following, multiple testing correction in our *F*_*ST*_ outlier analysis (*T*_*F-LK*_ = 16.2, *p*-value = 0.006, *q*-value = 0.52). Similarly, the extensively studied threonine/lysine polymorphism (Kreitman 1983; Powell 1997) that encodes the Fast and Slow allozyme variants at Alchohol dehydrogenase (Adh, 2L: 14617051) is not significantly differentiated following multiple testing correction in our *F*_*ST*_ outlier analysis (*T*_*F-LK*_ = 17.3, *p*-value = 0.004, *q*-value = 0.45).

**Figure 5.**
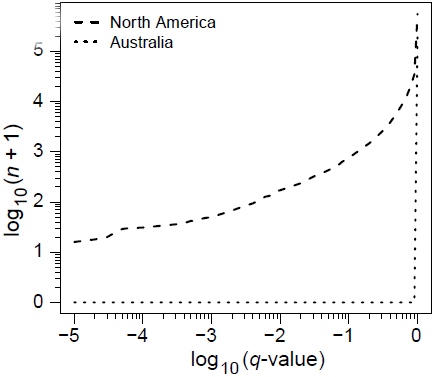
Number of significantly differentiated SNPs in North America and Australia at various FDR (*q*-value) thresholds.

While a limited number of polymorphisms were identified as significantly differentiated among North American populations, no significantly differentiated SNPs were observed among the Australian populations after correcting for multiple testing (Figure 5). Note that the genome-wide average *F*_*ST*_ among North American populations is lower than among Australian populations (0.025 vs. 0.08 respectively), suggesting that the lack of significantly elevated *F*_*ST*_ values in Australia is not due to a lack of population differentiation but rather a high genome-wide differentiation likely caused by recent secondary contact.

The exact number of SNPs with significantly elevated *F*_*ST*_ in any particular continent will be subject to a various of considerations including the number of sampled populations, the precision of allele frequency estimates, and the power of particular analytic methods to detect outlier *F*_*ST*_. Some of these factors vary between our North American and Australian samples and thus our power to detect significant elevation of *F*_*ST*_ will vary between continents. Therefore, we investigated the general patterns of differentiation and parallelism between the sets of populations sampled in North America and Australia. In addition, we also examined patterns of differentiation and parallelism between these continents and populations sampled from the Old-World (i.e., Europe and Africa).

For these analyses, we first examined whether SNPs that were highly differentiated among one set of populations were also differentiated in another set (hereafter, ‘co-differentiated’). To perform this analysis, we calculated the odds ratio (see *Materials and Methods*) that SNPs fell above a particular quantile threshold of the *F*_*ST*_ distribution in any two sets of populations (Figure 6A). We performed this analysis for SNPs that fell either within or outside of the large cosmopolitan inversions. We find that SNPs that are highly differentiated in North America are also highly differentiated in Australia. In addition, we find that SNPs that are highly differentiated in either North America or Australia are also highly differentiated between Europe and Africa. Although patterns of co-differentiation are higher among SNPs within the large, cosmopolitan inversion than for SNPs outside the inversions, the qualitative patterns remain the same for either SNP class suggesting that clinal variation in inversions *per se* does not drive the observed high levels of co-differentiation.

**Figure 6.**
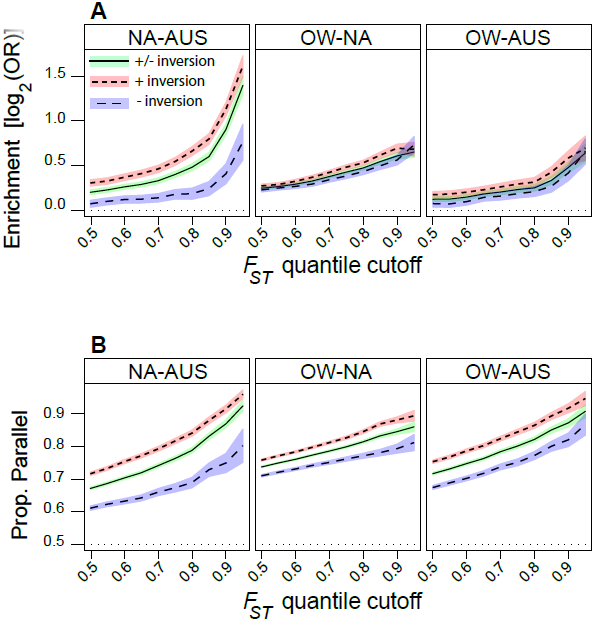
Patterns of co-differentiation and parallelism between North American, Australian, and Old-world populations. (A) log_2_ odds-ratio that SNPs fall above the *F*_*ST*_ quantile cut-off (x-axis) in both sets of populations (NA: North America; AUS: Australia; OW: Old-World). (B) Proportion of SNPs that vary in a parallel way given that they fall above the *F*_*ST*_ quantile cut-off in both sets of populations. Confidence bands represent 95% confidence intervals.

SNPs that are co-differentiated among temperate and tropical populations in North America, Australia, or the Old-World can be differentiated in a parallel way or at random among each geographic region. We show here that there is a high degree of parallelism at the SNP level, genome-wide, among polymorphisms that are highly differentiated in any two sets of populations (Figure 6B). Patterns of parallelism at highly co-differentiated SNPs are similar among SNPs within or outside the large cosmopolitan inversions again suggesting that clinal variation in inversions are not driving genome-wide patterns of parallelism.

High rates of co-differentiation and parallelism among temperate and tropical populations sampled throughout the world can be interpreted in two ways. On the one hand, these patterns could be taken as evidence of parallel adaptation to aspects of temperate environments. On the other hand, these patterns are consistent with the model presented above that North American and Australian populations are the result of recent secondary contact between European and African lineages of flies (see *Results: Genomic signals of secondary contact*).

To differentiate these alternative interpretations, we estimated rates enrichment of highly co-differentiated SNPs and rates of parallelism at highly co-differentiated SNPs among classes of polymorphisms that that we expect, *a priori*, to be more or less likely to contribute to local adaptation. We reasoned that SNPs falling in short-introns, which have been previously shown to evolve neutrally (Lawrie et al. 2013), would be the least likely to contribute to local adaptation. In contrast, SNPs in other functional classes (e.g., coding, UTR, intron) might be more likely to contribute to local adaptation along latitudinal clines (Reinhardt et al. 2014). We contrasted rates of co-differentiation and parallelism at these putatively functional SNP classes with rates at the short-intron (hereafter ‘neutral’) SNPs and at control SNPs matched to each class by several important biological and experimental features. These comparisons also take into account the spatial distribution of SNPs along the chromosome (see *Materials and Methods*). We reasoned that if parallel adaptive processes have contributed to genome-wide signals of co-differentiation and parallelism in Australia and North America, (1) some functional SNP classes would show a higher rate of co-differentiation and parallelism than neutral SNPs, (2) functional SNPs would show a higher rate of co-differentiation and parallelism than their control SNPs, and (3) neutral SNPs would show a lower rate of co-differentiation and parallelism than their control SNPs.

We find little evidence that various functional classes show differences in rates of co-differentiation or parallelism than either neutral SNPs or their matched controls (Fig. 7AB). Moreover, neutral SNPs show similar rates of co-differentiation and parallelism as their matched controls (Fig. 7AB). There is suggestive evidence that SNPs falling in 5’ UTRs show greater of co-differentiation than expected by chance, but this comparison is not significant after correcting for multiple tests (see *F*_*ST*_ > 95% Fig. 7A; *p*_*naive*_ = 0.01; *p*_*corrected*_ = 0.24). Moreover, highly co-differentiated SNPs in 5’UTR are not more likely to be parallel than expected by chance (Fig. 7B), suggesting that the observed excess of co-differentiation may be a statistical artifact. All other tests of excess co-differentiation or parallelism at different SNP classes were not significantly different from expectation (*p* > 0.05).

**Figure 7.**
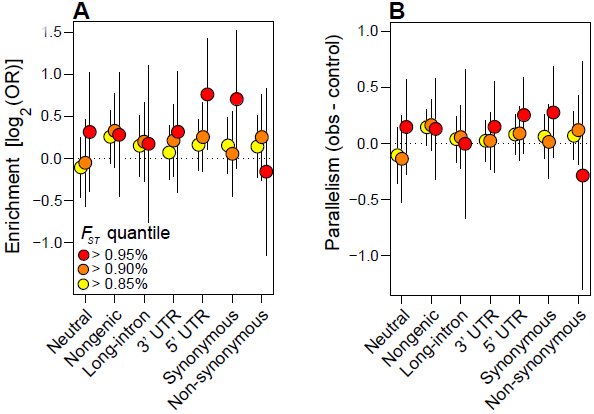
Patterns of (A) co-differentiation and (B) parallelism among various classes of SNPs relative to their matched controls. Vertical lines represent 95% confidence intervals. Horizontal dotted lines represents the null expectations. See *Materials and Methods* for details.

Taken together, the tests we performed to identify strong genomic signals of parallel adaptation along latitudinal clines were equivocal. We show that there were a modest number of *F*_*ST*_ outliers among North American populations sampled along a broad latitudinal cline and no observable *F*_*ST*_ outliers among Australian populations (Figure 5) suggesting that the bulk of the *F*_*ST*_ distribution is generated by the demographic history of this species. We show that SNPs with high *F*_*ST*_ among any one set of populations are likely to have high *F*_*ST*_ among other sets of population (Figure 6A). Furthermore, SNPs that are highly co-differentiated are likely to vary in a parallel fashion among geographic regions (Figure 6B). While this result could suggest parallel adaptation, it is also consistent with the dual colonization model we present above. Finally, we show that rates of co-differentiation and parallelism at highly co-differentiated SNPs are similar between functional SNPs, neutral SNPs, and their matched control SNPs (Figure 7) suggesting that the evolutionary forces shaping allele frequencies along latitudinal clines are similar across SNPs that are more-or less-likely to contribute to local adaptation.

## Discussion

Herein we report results from a series of analyses that (1) examine whether populations of *D. melanogaster* sampled throughout North America and Australia show signatures of recent secondary contact between European and African lineages, and (2) examine whether there is a genomic signal of spatially varying selection acting along latitudinal gradients. We find that both North America and Australia show several signatures of secondary contact (Figures 2-4). Notably, high latitude populations are closely related to European populations, whereas low latitude populations are more closely related to African ones. This result implies that a large portion of clinal variation within these continents could, in principal, be generated by the dual colonization of both North America and Australia (Figure 1). Consistent with this view, SNPs that are highly differentiated between temperate and tropical locales in either North America or Australia are also highly likely to be differentiated in a parallel way between Europe and Africa (Figures 6, 7). In addition, we report that genome-wide scans for significantly differentiated polymorphisms identified a limited number of outlier loci (Figure 5). Taken together, our results support the model that recent secondary contact in North America and Australia has generated clinal variation at a large fraction of polymorphisms genome-wide and that spatially varying selection acting at a moderate number of loci acts to slow the rate of genomic homogenization between geographically separated populations.

### Secondary contact and the generation of clinal variation in allele frequencies

Recent secondary contact between formerly (semi-) isolated populations is a potent force that can generate clinal variation genome-wide (Endler 1993). In *D. melanogaster*, high levels of genetic differentiation have been observed between temperate and tropical populations sampled in North America and Australia (Turner et al. 2008; Kolaczkowski et al. 2011; Fabian et al. 2012; Reinhardt et al. 2014). In North America at least, most of these highly differentiated SNPs vary clinally (i.e., in a roughly monotonic fashion along latitudinal gradients at false-discovery rate < 10%; Bergland et al. 2014). Moreover, surveys of allele frequencies along latitudinal clines in both North America and Australia at allozymes (Sezgin et al. 2004), SNPs (Sezgin et al. 2004; Lavington et al. 2014; Bergland et al. 2014), microsatellites (Gockel et al. 2001), and transposable elements (González et al. 2010) have repeatedly demonstrated that approximately one third of all surveyed polymorphisms are clinal in either continent. At face value the high proportion of clinal polymorphisms throughout *D. melanogaster*’s genome suggests that demographic processes such as secondary contact have contributed to the generation of clinal variation in this species among recently colonized locales (Bock & Parsons 1981; Keller 2007).

Accordingly, we tested if newly derived populations of *D. melanogaster* show signatures of recent secondary contact. Using a variety of tests, we show that genome-wide patterns of genetic variation from populations sampled in North America and Australia are consistent with recent secondary contact (Figures 2-4). While historical records from North America (Keller 2007) and Australia (Bock & Parsons 1981) suggest a single point of colonization of *D. melanogaster,* results from morphological, behavioral, and genetic studies reported here and elsewhere (Caracristi & Schlötterer 2003; Rouault *et al.* 2004; Duchen *et al.* 2013; Kao *et al.* 2014) suggest that a dual colonization scenario is more likely. At least for the Americas active trade between Europe and western Africa supports the model that North America represents a secondary contact zone.

Australia did not experience the same types of trade with the Old World and throughout the 19^th^ century intercontinental travel to Australia was primarily restricted to British ships. However, British ships traveling to Australia ported in South Africa and India then, after the opening of the Suez Canal in East Africa (Bach 1976). This raises the possibility that secondary contact between European and African fruit fly lineages could have occurred immediately prior to the successful colonization of Australia by *D. melanogaster* in the mid 19^th^ century (Bock & Parsons 1981). Under this mixed-lineage, single colonization scenario, rapid ecological sorting of colonizing lineages to temperate and tropical niches (Agosta & Klemens 2008) may have created a gradient where European flies were initially predominant at high latitudes and African flies predominant at low latitudes within Australia.

Although secondary contact is capable of generating patterns of clinal variation genome-wide, clines generated through this demographic process are transient. As admixed populations approach migration-selection equilibrium, clines at neutral loci should attenuate. Moreover, once at equilibrium, neutral differentiation should be minimal (Slatkin 1987) for species such as *D. melanogaster* where *Nm* has been estimated to be on the order of ∼1 (Yamazaki *et al.* 1986; Singh & Rhomberg 1987) and long-distance dispersal is believed to be frequent (Coyne & Milstead 1987).

Thus, the critical question in determining whether the vast amount of clinal variation in North American and Australian flies has been generated by demography or selection is whether or not this species is at migration-selection equilibrium in these continents. There are several reasons why we suspect this species is not at equilibrium. First, *D. melanogaster* appeared in North America and Australia in the mid-to late 19^th^ century (Bock & Parsons 1981; Keller 2007), or on the order of 1000 generations ago, assuming approximately 10 generations per year. Estimates of local, demic *N* are on the order of 10^4^ (McKenzie 1980; McInnis et al. 1982; Powell 1997) implying that *m* is on the order of 10^-4^ (if *Nm* ∼ 1). If these estimates are accurate to the order of magnitude, it would take approximately 2500 generations to get about half way to equilibrium (Whitlock 1992) or ∼10,000 generations to fully approach equilibrium (Whitlock & McCauley 1999). Thus, from a simple demographic perspective, it would seem unlikely that *D. melanogaster* has reached migration-selection-drift equilibrium.

Others have suggested that non-African populations of *D. melanogaster* are not at equilibrium. In general, non-African populations of *D. melanogaster* show a reduction in diversity coupled with an excess of rare variants (Mackay et al. 2012). This genome-wide pattern is consistent with a population bottleneck during colonization followed by population expansion. Others have noted that non-African populations of *D. melanogaster* also have higher levels of linkage-disequilibrium (LD) than expected under the standard neutral model (Andolfatto & Przeworski 2000; Haddrill *et al.* 2005; Langley *et al.* 2012) whereas LD in African populations is more consistent with neutrality (Andolfatto & Wall 2003 *cf.* Langley et al. 2012). Although genome-wide elevation of LD could be caused by various factors including pervasive positive-or negative-selection, admixture would also possibly generate this signal.

Previous studies examining departure from equilibrium models in *D. melanogaster* have concluded that caution should be taken when conducting genome-wide scans for positive-selection given the non-equilibrium nature of this species (Andolfatto & Przeworski 2000). Notably, demographic forces such as population bottlenecks can, in principal, mimic many of the signatures left by some types of adaptive evolution. A complimentary approach to quantify the magnitude of adaptive evolution and to identify loci subject to selection is to identify polymorphisms that are differentiated between populations that are subject to divergent selection pressures. However, results presented here demonstrate that, for *D. melanogaster* at least, signatures of adaptive evolution from genome-wide patterns of differentiation along latitudinal clines in newly derived populations in North America and Australia should be taken with a similar or even greater degree of caution as traditional scans for recent, positive selection.

### Spatially varying selection and the maintenance of clinal variation in allele frequencies

Whereas secondary contact is capable of generating clinal variation, spatially varying selection is required for its long-term maintenance. There is little doubt that populations of *D. melanogaster* living along broad latitudinal clines in temperate environments have adapted to spatially varying selection pressures. Support for the idea of local adaptation along latitudinal clines comes from three main lines of evidence.

First, certain phenotypes show repeatable clines along latitudinal and altitudinal gradients that mirror deeper phylogenetic variation among temperate and tropical species. For instance, aspects of body size vary clinaly in North America (Coyne & Beecham 1987) and Australia (Kennington et al. 2003) as well as along altitudinal/latitudinal clines in India (Bhan et al. 2014) and altitudinal clines within Africa (Pitchers et al. 2013; Klepsatel et al. 2014). Given such patterns of parallelism within and among continents, including within the ancestral African range, the most plausible explanation is that parallel selection pressures have generated these patterns of latitudinal and altitudinal variation. These intraspecific clines mimic interspecific patterns among temperate and tropical endemic drosophilids following Bergmann’s rule (Blanckenhorn & Demont 2004; Shelomi 2012) again implicating that natural selection has shaped these patterns of genetically based, phenotypic variation.

Second, certain genetic and phenotypic clines in *D. melanogaster* have shifted over decadal scales. Shifts in these clines are consistent with adaptation to aspects of global climate change wherein alleles common in low-latitude populations have become more prevalent in high-latitude ones over the last 20 years (Hoffmann & Weeks 2007).

Finally, here we identify several hundred polymorphisms in North America that are significantly differentiated (see *Results*, Figure 5, and Supplemental Table 3). Although the function of many of these polymorphisms is presently unknown, several are within the genes known to affect life-history traits and correlates that vary among temperate and tropical populations (see *Results*).

The identification of significantly differentiated SNPs within North America can be taken as evidence of local adaptation to spatially varying selection pressures. However, the observation that two SNPs (one in *cpo* and one in *Adh*) that each likely contribute to local adaptation fall in an upper, but not extreme, tail of the *F*_*ST*_ distribution suggests that there are many more ecologically relevant and functional polymorphisms that have contributed to local adaptation in *D. melanogaster.* However, the signal of high differentiation caused by spatially varying selection at these SNPs is likely masked by recent admixture that has contributed to a high level of differentiation genome-wide. In light of these results, we suggest that scans for local adaptation based on patterns of genetic differentiation in *D. melanogaster* are an important first step in identifying adaptively differentiated clinal polymorphisms but that additional evidence, such as functional validation (Schmidt *et al.* 2008; Paaby *et al.* 2010; 2014), should be gathered before concluding that differentiation is caused by adaptive processes.

### Conclusions

It has long been recognized that genetic differentiation among populations can be caused by both adaptive and demographic (neutral) processes (Wright 1943). Due to *D. melanogaster*’s large effective population size (Karasov et al. 2010), high migration rate (Coyne & Milstead 1987), and rapid decay of linkage disequilibrim (Mackay et al. 2012) others have concluded that differentiation among populations sampled along latitudinal gradients is primarily caused by spatially varying selection. Work presented here supports the notion that spatially variable selection does contribute to some differentiation among populations.

However, several genome-wide signatures presented here (Figures 2-4) and elsewhere (Caracristi & Schlötterer 2003; Duchen *et al.* 2013; Kao *et al.* 2014; Bergland *et al.* 2014) indicate that populations of flies in North America and Australia result from admixture of European and African lineages. High-latitude (temperate) populations in North America and Australia are more closely related to European populations whereas low-latitude (tropical) populations are more closely related to African ones (Figures 2, 3) suggesting that admixture occurred along a latitudinal gradient and that this demographic event generated clinal genetic variation at roughly 1/3 of all common SNPs (Bergland et al. 2014). These colonizing lineages of flies were likely already differentially adapted to the temperate and tropical conditions that they encountered in North America and Australia. Consequently the recent demographic history of this species in North America and Australia is collinear with both local adaptation within these newly colonized continents and among the ancestral ranges. One practical consequence of the collinearity of demography and adaptation is that the identification of clinality at any particular locus cannot be taken exclusively as evidence of spatially varying selection.

The collinearity of demography and adaptation may be a general feature of a variety of species. For instance, the successful colonization of novel locales by invasive species such as *D. melanogaster* is often facilitated by multiple waves of invasion. In addition to increasing propagule pressure (Simberloff 2009), multiple independent invasions are also thought to facilitate invasion success by buffering the loss of genetic diversity that accompanies bottlenecks associated with colonization. Moreover, some evidence suggests that rapid adaptation following invasion (reviewed in Dormontt et al. 2011), often results directly from hybridization between independently invading subpopulations (reviewed in Lee 2002). If invasive species are widely distributed in their native range (Bates et al. 2013), this raises the possibility that successful invasions may often result from admixture of populations that are already differentially adapted to selection pressures that vary along broad spatial gradients in the introduced ranges.

The collinearity of demography and adaptation may also occur in temperate endemic species. This phenomenon may be particularly true in marine taxa as a result of admixture between high-latitude refugial populations and low-latitude populations following the last glacial maxima ∼20,000 yeas ago (Bernatchez & Wilson 1998; Maggs *et al.* 2008). For these species, secondary contact between high-and low-latitude populations would result in clinal variation at neutral loci as well as loci that contribute to adaptation along latitudinal gradients (e.g., Adams et al. 2006). In this scenario, neutral and adaptive genetic clines may be indistinguishable.

The ability to identify targets of spatially varying selection is now possible in many model and non-model species (Li et al. 2008; Savolainen et al. 2013). While these genome-wide approaches can be extremely powerful, great care must be taken to ensure that signals of local adaptation are uniquely identifiable and independent from pervasive (but often subtle) signals of demography. *F*_*ST*_ outlier approaches can in some cases enable the detection of loci that underlie local adaptation following complex and confounding demographic scenarios. However, as we show here, loci known to underlie functionally relevant, phenotypic variation are not necessarily detected as statistically significant outliers, likely due to the pervasive signals of recent demographic events. To ameliorate these concerns, we suggest that proposed genetic targets of spatially varying selection be functionally verified, or that patterns of spatial variation at loci known *a priori* to underlie fitness related phenotypic variation be investigated. Alternatively, the genomic targets of local adaptation can be identified by examining population differentiation over small spatial scales (Richardson et al. 2014) or over short time periods (Bergland et al. 2014) that are likely to be orthogonal to the demographic history of the focal species.

## Acknowledgements

We thank Joyce Kao, Heather Machado, and Annalise Paaby for insightful comments on earlier versions of this manuscript. We thank David Lawrie for providing lists of neutrally evolving SNPs and Ary Hoffmann for graciously providing isofemale lines from Australia. Finally, we thank Richard Hudson for help with *ms* and Nick Patterson and Peter Ralph for assistance interpreting *D* statistics. AOB was supported by an NIH National Service Research Award (F32 GM097837). JG is a Ramon y Cajal fellow (RYC-2010-07306) supported by grants from the European Commission (Marie Curie CIG PCIG-GA-2011-293860) and from the Spanish Government (Fundamental Research Projects Grant BFU-2011-24397). This work was supported by NSF DEB 0921307 (awarded to PS) and NIH R01GM089926 (awarded to PS and DP).

## Data Accessibility

- Raw DNA sequence data for Inisfail and Yering Station samples: NCBI SRA: BioProject 270869
- *ms* code to generate simulations of various colonization models: Data Dryad: doi: 10.5061/dryad.gg5nv
- All other genomic data are previously published.

## Author Contributions

AOB and RT analyzed the data. JG generated sequencing libraries. AOB, RT, JG, PS, and DP wrote the manuscript.

---

**Supplemental Text 1**. Neighbor-joining trees for each chromosome in Newick format.

**Supplemental Figure 1.**
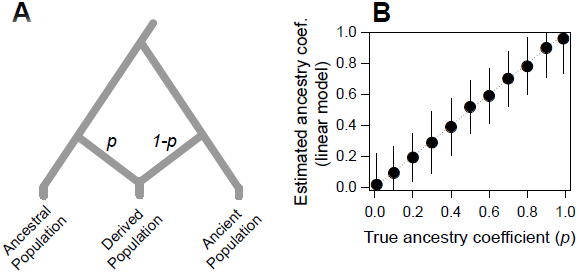
Verification that the linear model method of ancestry proportion estimation is accurate.

**Supplemental Figure 2.**
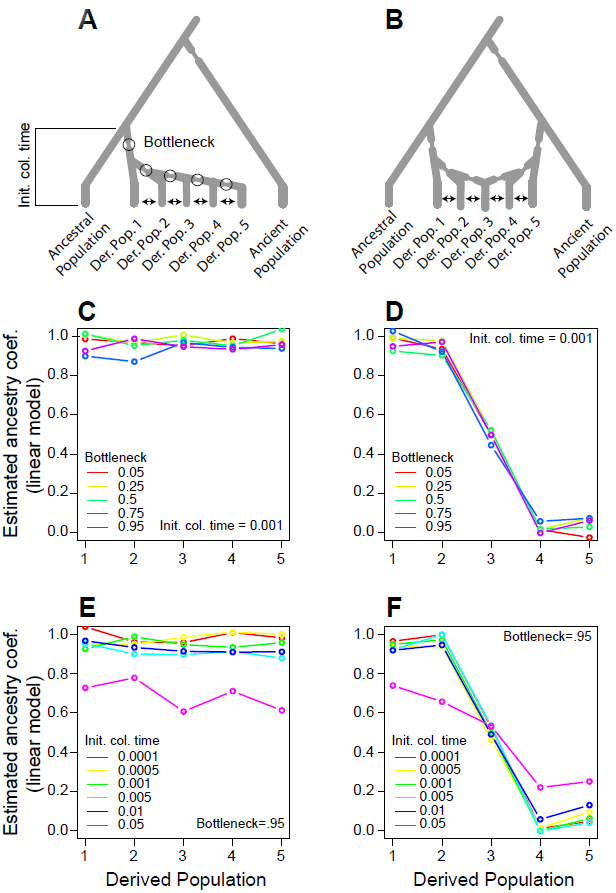
Verification that the linear model method of ancestry proportion estimation is robust under various demographic scenarios.

**Supplemental Figure 3.**
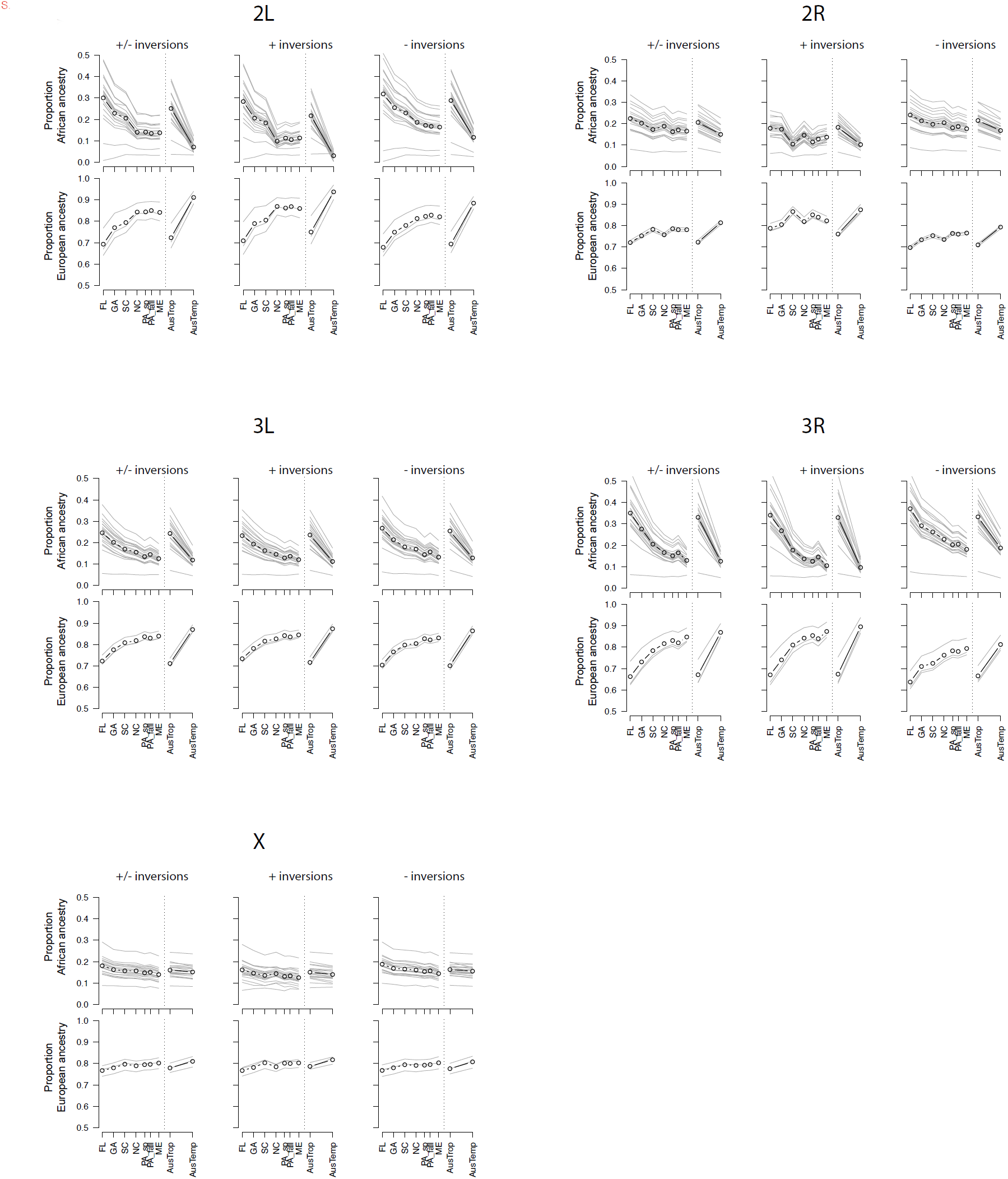
Ancestry estimates for each chromosome partitioned by inversion status.

**Supplemental Table 1**. *f*_3_ and *D* statistics for each pair-wise combination of African and European populations.

**Supplemental Table 2**. Annotated table of F_ST_ outlier SNPs at *q* < 0.15 in North America.

